# Phenotype prediction from genome-wide genotyping data: a crowdsourcing experiment

**DOI:** 10.1101/2020.08.25.265900

**Authors:** Olivier Naret, David AA Baranger, Sharada Prasanna Mohanty, Bastian Greshake Tzovaras, Marcel Salathé, Jacques Fellay, with the openSNP and crowdAI community

**Affiliations:** School of Life Sciences, Ecole Polytechnique Fédérale de Lausanne, Switzerland; Department of Psychological and Brain Sciences, Washington University, St. Louis, MO, USA; Lawrence Berkeley National Laboratory, Berkeley, CA, USA; Department for Applied Bioinformatics, Goethe University, Frankfurt am Main, Germany; Center for Research and Interdisciplinarity (CRI), Université de Paris, INSERM U1284, Paris, France

**Keywords:** crowdsourcing, genomic prediction, open science, openSNP, polygenic risk score, machine learning

## Abstract

**Background:** The increasing statistical power of genome-wide association studies is fostering the development of precision medicine through genomic predictions of complex traits. Nevertheless, it has been shown that the results remain relatively modest. A reason might be the nature of the methods typically used to construct genomic predictions. Recent machine learning techniques have properties that could help to capture the architecture of complex traits better and improve genomic prediction accuracy.

**Methods:** We relied on crowd-sourcing to efficiently compare multiple genomic prediction methods. This represents an innovative approach in the genomic field because of the privacy concerns linked to human genetic data. There are two crowd-sourcing elements building our study. First, we constructed a dataset from openSNP (opensnp.org), an open repository where people voluntarily share their genotyping data and phenotypic information in an effort to participate in open science. To leverage this resource we release the ‘openSNP Cohort Maker’, a tool that builds a homogeneous and up-to-date cohort based on the data available on opensnp.org. Second, we organized an open online challenge on the CrowdAI platform (crowdai.org) aiming at predicting height from genome-wide genotyping data.

**Results:** The ‘openSNP Height Prediction’ challenge lasted for three months. A total of 138 challengers contributed to 1275 submissions. The winner computed a polygenic risk score using the publicly available summary statistics of the GIANT study to achieve the best result (*r*^2^ = 0.53 versus *r*^2^ = 0.49 for the second-best).

**Conclusion:** We report here the first crowd-sourced challenge on publicly available genome-wide genotyping data. We also deliver the ‘openSNP Cohort Maker’ that will allow people to make use of the data available on opensnp.org.

## Background

As costs for genetic analyses keep dropping, genetic testing is becoming more available and affordable for increasing numbers of people - a trend that can be seen in the rising number of customers that use Direct-To-Consumer (DTC) genetic testing services like 23andMe and AncestryDNA [1]. Decreasing costs and increased availability have lead to the creation of a number of genomic data resources, such as the Personal Genome Project [2], DNA.land [3] and openSNP [4]. Amongst these data resources, openSNP is unique, in that it offers open participation and open access to the data: Participants of openSNP can use the platform to openly share their existing DTC genetic test data, putting their data in the public domain. In addition, participants can share phenotypic traits, such as eye color, hair color or height. Since its start in 2011, over 5,000 people have used the platform to make their genetic data available.

Crowd-sourced competitions of data analysis have become more and more popular in the past few years allowing data science experts and enthusiasts to collaboratively solve real-world problems, through online challenges. This approach allows the broad exploration of the model space on a specific dataset by people with data analysis skills coming from very different backgrounds. In the context of genomic prediction of complex diseases, it is unprecedented. While the most widely used platform, kaggle.com, offers monetary rewards, crowdai.org is more academic-centered and offered the winner the opportunity to present her work at a scientific conference.

We hereby present a crowd-sourcing experiment where participants could compete on crowdai.org to produce the best possible prediction of the height phenotype using data from opensnp.org.

## Materials and methods

### openSNP Cohort Maker

Because on opensnp.org, no restrictions are enforced on what users can upload, after downloading the data dump of the whole community, there is a need for in-depth data curation to produce a clean cohort of genome-wide genotyped individuals. To make these data accessible to anyone, we developed the openSNP Cohort Maker tool that through a systematic approach produced a clean and up-to-date openSNP cohort of genome-wide genotyped individuals.

When running the openSNP Cohort Maker, the data processing starts by downloading the archive containing all data that were uploaded on opensnp.org by the community. Then, files are removed if: they are not text or compressed text; they correspond to exome sequencing; they are genotyping data from decodeme; they are set on the GRCh38 reference; they are corrupted. For individuals who submitted multiple genome-wide genotyping data, either as duplicates or from different DTC companies, only the largest file is kept. A set of tools are integrated into the pipeline: genotyping data with coordinates based on NCBI36 are upgraded to match the GRCh37 reference [5] with liftOver [6]; PLINK [7–9] is used to convert file formats. VCFtools [10] is used to sort variants; BCFtools [11] is used to normalize reference and alternate alleles on the GRCh37 reference genome, rename samples, index files, and finally merge all individuals into one file. The output file can be directly imputed on the Sanger Imputation Service [12]. The openSNP Cohort Maker is available on GitHub S1. Software. Leveraging parallel computing, with 28 CPUs it takes 16 hours to produce a single file containing the curated openSNP cohort. From the initial archive containing 2487 different individuals, 2341 remain after filtering. From those, 2034 are from 23andme, 186 are from ancestry.com, and 121 are from ftdna-illumina.

### CrowdAI Challenge

The dataset that we used for the challenge was produced by the openSNP Cohort Maker and imputed on Sanger Imputation Service with HRC (r1.1). We sent to the opensnp.org community a survey asking for their height, allowing us to create a dataset regrouping 921 individuals with both height phenotype and genotyping data.

Challenge participants could use two versions of the genotyping data. One version was a sub-dataset containing 9,894 genetic variants, including the top 9,207 variants (*p* < 5×10^−3^) associated with height in the GIANT study [13], and 687 Y chromosome variants. The second versions was a full dataset containing 7,252,636 variants which passed a quality threshold, defined as an imputation score *INFO* > 0.8, genotyping missingness frequency *F*_*m*_ < 0.1, and a Hardy-Weinberg equilibrium exact test *p*− *value* < 1*e*^−50^. Both versions of the data were given in the VCF format, as well as in an additive format where each genetic variant is represented by 0 (homozygous for reference), 1 (heterozygous), 2 (homozygous for the alternate allele) or NA (missing data or variants of allosomes), easier to handle for participants unfamiliar with genetics.

The data were separated into two sets, a training set with 784 samples and a test set of 137 samples (an 85/15 split). The challengers were provided the training set with the genotyping data and the height phenotype and the test set with the genotyping data only. The challengers needed to train their model on the training set and produce predictions for the samples of the test set. The test set predictions were then submitted to the CrowdAI platform for evaluation and scoring. The score was produced based on the Pearson’s correlation (*r*^2^) between the predicted and true height. The challengers could submit as many prediction models as they wanted in an attempt to improve their method and beat their best score. The scoring method was protected from known exploits [14]. The data are available online on the zenodo platform S1. Dataset, and the webpages presenting the challenge S1. Appendix and the leaderboard S2. Appendix have been saved to PDF from the CrowdAI platform.

## Results

A total of 138 challengers participated, contributing a total of 1275 submissions. The winner computed a polygenic risk score (PRS) using the publicly available summary statistics of the GIANT study to achieve the best result (*r*^2^ = 0.53 versus *r*^2^ = 0.49 for the second-best).

The winning method was based on PRS. The training set and testing set were combined for quality control and data preparation. As self-reported sex was not provided, participant’s chromosomal sex (i.e. XX vs XY) was imputed using PLINK, which uses the X chromosome inbreeding coefficient (F) to impute sex. Standard cutoffs were used, whereby *F* < 0.2 yielded an XX call, while *F* > 0.8 yielded an XY call. One participant yielded an F of exactly 0.2, and was removed from subsequent analyses (they were in the training data). Of the remaining 920 individuals, 396 (43%) were XX, and 524 (57%) were XY.

The openSNP platform contains genomic data of relatives. The presence of relatives has the potential to bias results, as closely-related individuals will dominate the estimation of principal components and will inflate prediction accuracy statistics [15]. The genetic relationship between participants was calculated using the PLINK computation of identity-by-descent (IBD), which is an estimate of the percent of the genome (excluding sex chromosomes) shared between two individuals. The IBD analysis identified seven pairs of strongly-related individuals (first-cousin or greater), including two pairs of monozygotic twins. The analysis also identified a surprising cluster of 18 individuals estimated to be 3^*rd*^ cousins, or equivalent. All but one member of each family-group was removed from analyses (N=24), all from the training data. It is well-established that the frequency of genetic variants and correlational structure of the genome differs across ancestral populations [15–17]. These differences are the major barrier to combining genomic data across ancestries in genome-wide association studies [18, 19]. Genome-wide principal components were computed using PLINK. A scree plot of eigenvalues indicated an elbow at three components. The large eigenvalue of the first principal component, and the shape of all three components, clearly showed that both the training and testing data contained participants of multiple ancestries (i.e. participants of European, African, and Asian ancestry were present in both data sets), though the majority of participants were of European descent.

Genomic data were further processed in PLINK, following the steps outlined by PRSice [20] for the computation of PRS. This included removal of variants that were missing for more than 2% of participants, removal of variants with a minor-allele-frequency less than 0.02, removal of variants with a Hardy-Weinberg equilibrium exact test *p*_*value*_ < 10^−6^, removal of variants within the major histocompatibility complex on chromosome 6, and removal of non-synonymous variants. Because neighboring genetic variants can be correlated due to linkage disequilibrium, genetic variants were clumped in PLINK, wherein groups of variants correlated at *r*^2^ > 0.1 were identified across a sliding window of 250 kilobases. Within each group, only the variant with the lowest p-value in the GIANT genome-wide association study of height [13] was retained.

A PRS is a metric reflecting an individual’s genetic burden for a disease or trait of interest. [21, 22]. Prior work on the genetic basis of height has found that a PRS for height captures over 20% of the variance in independent samples [13]. PRS are calculated by averaging the number of disease-associated alleles, weighted by their effect size, from an independent study [23]. Put differently, a linear regression predicting the outcome trait is modeled at each individual variant, using the effect size from an independent study. These predictions are then averaged across all models. The one free parameter is the decision of which variants to include in the calculation of the PRS. Typically, the significance of the association of each variant in the independent study is used. Thus, multiple scores are calculated, including only variants that are associated below different p-value thresholds (e.g. *p* < 5 ×10^−1^, *p* < 5 ×10^−2^, *p* < 5 ×10^−3^, etc.). While a PRS including all variants (i.e. *p* < 1.0) typically does not perform the best, neither does a PRS including only variants which surpass family-wise error rate correction for multiple comparison (i.e. *p* < 5 × 10^−8^). Finally, the confounding effects of ancestral populations apply to PRS analyses as well. PRS work best when the independent study (e.g. the GIANT study used here) was conducted within a homogeneous sample of participants all of whom are from the same ancestral background, and when the sample the PRS is being calculated for is of the same background. For instance, PRS computed from studies of individuals of European descent are well-known to produce biased results in samples of African or Asian descent [24, 25].

PRSice and PLINK were used to compute PRS for height in the openSNP sample, using the results from the GIANT study of height. PRS were computed at 14 different p-value thresholds (*p* < 10^−8^ to *p* < 1.0), shown in Fig 1. Linear regressions predicting height in the training data were fit in R [26]. Chromosomal sex was the first variable included in the model, followed by the top three genome-wide principal components, which help to control for differences in ancestral background [27]. Chromosomal sex predicted 46.81% of the variance in the training data, and the addition of the three principal components subsequently explained 0.91% of variance. Finally, each of the 14 PRS were added to the model and compared. The PRS at *p* < 1*x*10^−5^ was observed to perform best, and captured an additional 10.08% of variance. Thus, the final linear regression model explained a total of 57.80% of the variation of height in the training data set, shown in Fig 2. Predictions for height in the test data set were then generated from this regression model, predicting 53.45% of variance (MSE = 47.32).

**Fig 1.**
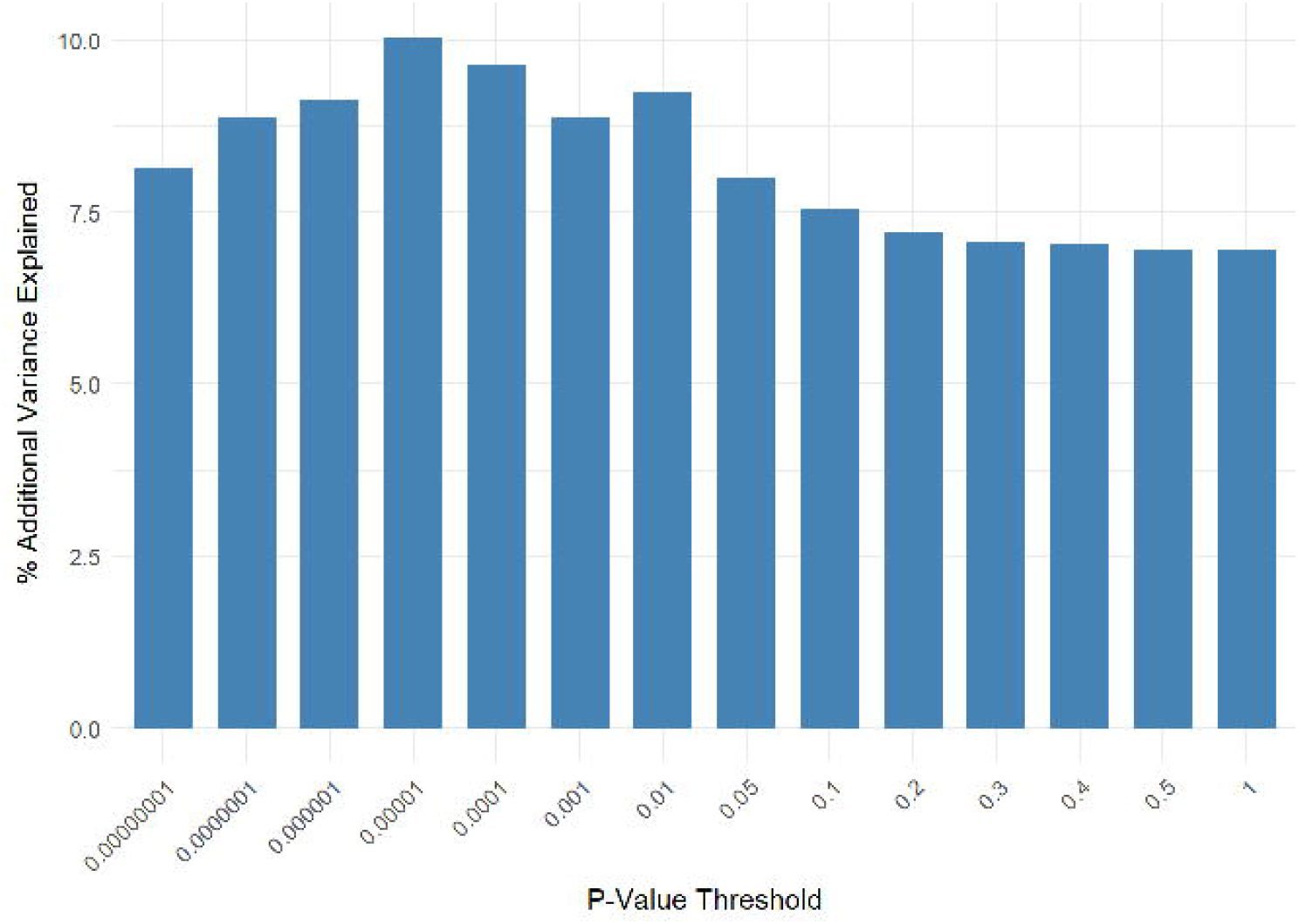
Variance explained as a function of *p*_*value*_ threshold. PRS were produced at a range of p-value thresholds (x-axis). Y-axis represents Nagelkerke’s r-squared from training-sample linear regressions. The model with the best performance in the training data (*p* < 5 × 10^−4^) was then used to predict height in the test-sample.

**Fig 2.**
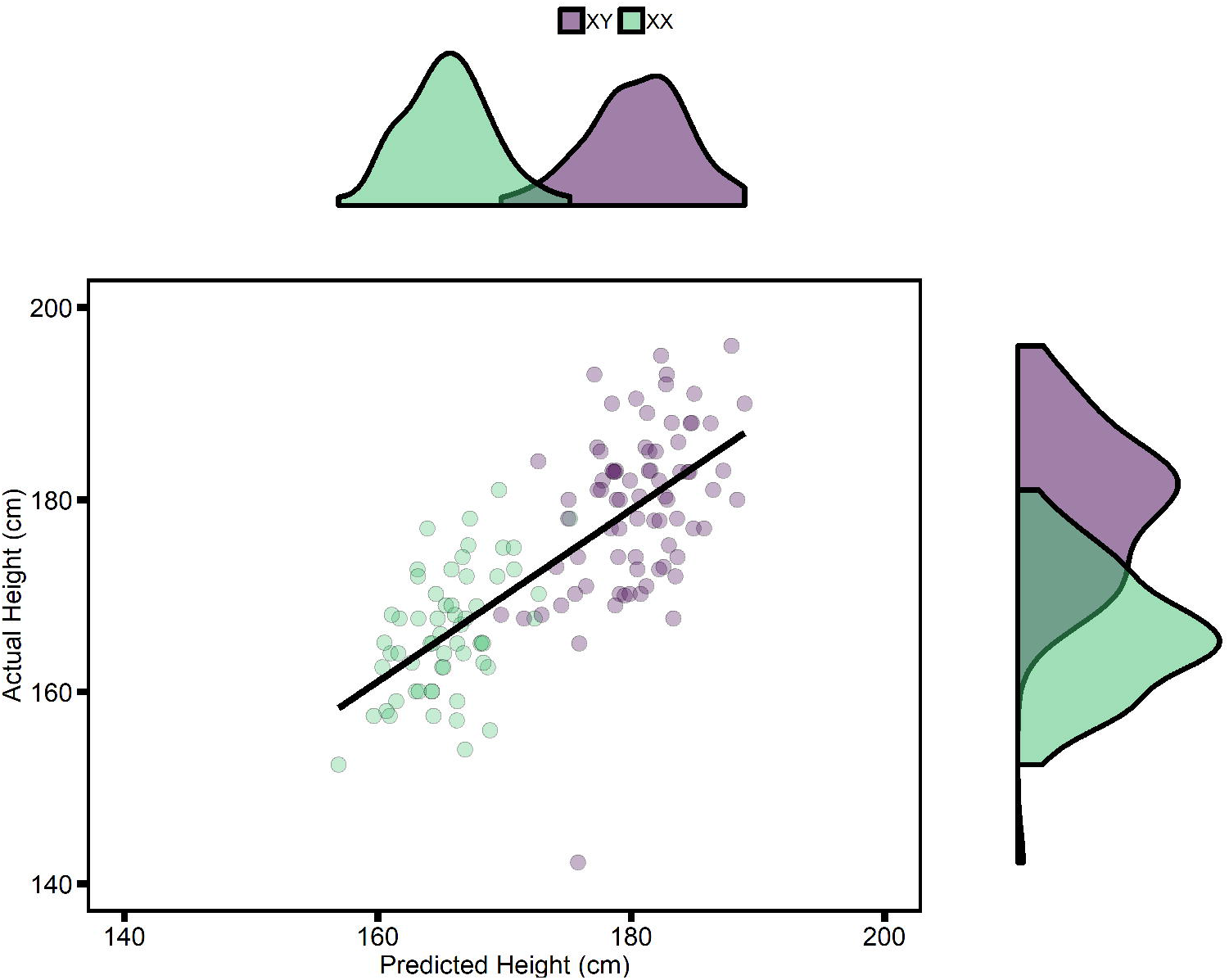
Predicted height distribution versus real height distribution. Predicted height in the training-sample (x-axis) is displayed relative to true height (y-axis). Points are colored by chromosomal sex. X and y-axis density plots show the predicted and true overlap of height between the sexes.

## Discussion

Height is an extremely polygenic trait where even the hundreds of genome-wide significant variants contribute all together for only a small portion of heritability [28]. Because of the modest size of the OpenSNP cohort, the lack of statistical power was the main difficulty for the challengers to capture the association signals coming from the genetic variants. The winning model of the challenge incorporated the GWAS summary statistics from the GIANT study to compute a PRS, in addition to deriving each participant’s sex. It should be noted that PRS is a standard and widely-used technique in the field of statistical genetics. While cross-population PRS have been shown to be unreliable in multiple cases, such as Type II Diabetes [29], coronary artery disease [30], and height [31], the similarities between the GIANT and openSNP cohorts were sufficient to provide a winning strategy. This is likely because only a small portion of samples were of non-European ancestry (7%).

So far, PRS are classically limited to additive models which might not represent the whole complexity of the genetic architecture of some traits. Indeed, the phenotypic variance explained by PRS remains modest in comparison to the heritability of the traits [32] (the so-called ‘missing heritability’ problem). The inability to consider gene-gene interactions is one of the many factors potentially explaining for this PRS weakness. In this case, the effect of a variant depends on the presence or absence of another variant, a mechanism that is not captured by additive models and accounts for an unknown part of the phenotypic variance [33]. Eventually, more advanced statistical approaches relying on machine learning could improve on the prediction accuracy provided by purely additive risk scores. Because of the diversity in available methods and the world-wide distribution of excellent data scientists, we believe that crowd-sourcing approaches represent a promising strategy to help improve phenotypic prediction from large-scale genomic data.

## Conclusion

Because of privacy concerns, studies relying on crowd-sourcing are almost impossible to set up in the field of human genomics. A first experiment was carried out in 2016 to predict anti-TNF treatment response in rheumatoid arthritis [34], but participants had to apply to participate in the challenge. Here - thanks to the OpenSNP community - we released the first crowd-sourced and fully open challenge based on publicly available genome-wide genotyping data. The competition attracted 138 challengers, with diverse backgrounds, from the vibrant machine learning community. It resulted in the assessment of a broad variety of methods for genotype-based phenotypic prediction through a total of 1275 submissions. We hereby also report a tool to create an up-to-date and curated OpenSNP cohort, making this open genomic resource much more user-friendly.

## Supporting information

**S1. Appendix Released challenge presentation**.

**S2. Appendix Challenge leaderboard**.

**S1. Software OpenSNP cohort maker:** https://github.com/onaret/opensnp-cohort-maker

**S1. Dataset Challenge dataset:** https://zenodo.org/record/1442755#.XlTwyHVKh1M

## Notes

https://zenodo.org/record/1442755#.XlTwyHVKh1M

### Competing Interest Statement

The authors have declared no competing interest.

